# Topologically associating domain boundaries are enriched in early firing origins and restrict replication fork progression

**DOI:** 10.1101/2020.10.21.348946

**Authors:** Emilia Puig Lombardi, Madalena Tarsounas

## Abstract

Topologically associating domains (TADs) are units of the genome architecture defined by binding sites for the CTCF transcription factor and cohesin-mediated loop extrusion. Genomic regions containing DNA replication initiation sites have been mapped in the proximity of TAD boundaries. However, the factors that determine this positioning have not been identified. Moreover, the impact of TADs on the directionality of replication fork progression remains unknown. Here we use EdU-seq technology to map origin firing sites at 10 kb resolution and to monitor replication fork progression after restart from hydroxyurea arrest. We show that origins firing in early/mid S-phase within TAD boundaries map to two distinct peaks flanking the centre of the boundary, which is occupied by CTCF and cohesin. When transcription is inhibited chemically or deregulated by oncogene overexpression, replication origins become repositioned to the centre of the TAD. Furthermore, we demonstrate the strikingly asymmetric fork progression initiating from origins located within TAD boundaries. Divergent CTCF binding sites and neighbouring TADs with different replication timing (RT) cause fork stalling in regions external to the TAD. Thus, our work assigns for the first time a role to transcription within TAD boundaries in promoting replication origin firing and demonstrates how genomic regions adjacent to the TAD boundaries could restrict replication progression.

## INTRODUCTION

In eukaryotic cells, thousands of specialized DNA regions, termed replication origins, need to be activated in order to replicate DNA accurately and efficiently before cell division (Leonard and Méchali 2013; Hyrien 2015). Over the past years, a comprehensive understanding of the mechanisms underlying the DNA replication process and the molecular components of the machinery needed for its completion has been acquired. During the G1 phase of the cell cycle, the assembly of the pre-replicative complex (pre-RC) licenses origins for potential activation in the subsequent S-phase (Siddiqui et al. 2013). Pre-RC assembly is completed in G1, by the loading of the replicative helicase complex MCM2-7 at each potential origin (Méndez and Stillman 2003; Li et al. 2015). Helicase loading requires the origin recognition complex (ORC), the cell division control protein 6 (CDC6) and the DNA replication licensing factor CDT1 to complete (Gambus et al. 2011). Upon entry into S-phase, the replicative helicase is activated by cyclin-dependent kinases (CDKs) and the DBF4-dependent kinase (DDK), which facilitate CDC45 and GINS recruitment to the origin and formation of the CMG (CDC45/MCM2-7/GINS) helicase complex (Moyer et al. 2006; Ilves et al. 2010). Active CMG initiates DNA unwinding and enables replisome assembly, whilst traveling ahead of the replication fork (reviewed in (O’Donnell et al. 2013; Riera et al. 2017)).

Next-generation sequencing (NGS)-based approaches led to genome-wide mapping of potential replication origins in the human genome. Three independent approaches, relying on the direct identification of DNA replication initiation intermediates, have been initially established: 1) SNS-seq, which relies on isolation of small nascent DNA strands enriched by lambda DNA exonuclease digestion, combined with NGS (Besnard et al. 2012; Picard et al. 2014; Cayrou et al. 2015) or microarrays (SNS-chip; (Cadoret et al. 2008; Sequeira-Mendes et al. 2009; Cayrou et al. 2012)); 2) Bubble-seq, which relies on capturing and sequencing restriction fragments that contain DNA replication initiation sites or “bubbles” (Mesner et al. 2013); and 3) OK-seq, which relies on sequencing purified Okazaki fragments for genomewide evaluation of replication fork polarity and mapping of broad (~30 Kb) initiation and termination zones (Petryk et al. 2016). More recently developed methods relying on high-throughput sequencing of newly-synthesized DNA (Langley et al. 2016; Macheret and Halazonetis 2018; Tubbs et al. 2018) yielded high-resolution maps of discrete origin firing sites within the genome. As a prominent example, the EdU-seq technology combines the incorporation of the thymidine analogue 5-ethynyl-2’-deoxyuridine (EdU) with NGS, allowing genome-wide mapping of replication origins at high resolution (1 to 10 Kb), as well as monitoring replication fork progression (Macheret and Halazonetis 2018; Macheret and Halazonetis 2019). EdU-seq has reported mapping of constitutive (CN) replication origins in three different cell lines (U2OS, HeLa and RPE1), with the majority of origins overlapping in the three cell lines (Macheret and Halazonetis 2018).

Although all potential replication origins are licensed, only 10% are activated in mammalian cells (Rivera-Mulia and Gilbert 2016). Additionally, those origins that do fire follow a defined temporal order defined as replication timing (RT; (Rhind and Gilbert 2013)). Synchronous replication firing mediated by clusters of replication initiation events is a feature of large chromosomal domains termed constant timing regions (CTRs), which are separated at reproducible locations by timing transition, defined as sites where RT transitions between early/mid/late S-phase (Hiratani et al. 2008; Pope et al. 2014; Marchal et al. 2019). Interestingly, CTRs consist of smaller units, or replication domains (Hiratani et al. 2008). Their boundaries overlap with experimentally mapped topologically associating domain (TAD) boundaries. TADs are three-dimensional structural subunits of the genome, mapped using high-resolution chromosome conformation capture (Hi-C). TADs are defined as self-interacting genomic regions, or regions within which sequences interact more frequently with each other compared to their interactions with sequences outside of the TAD (Dixon et al. 2012; Nora et al. 2012). TADs are enriched in histone modifications indicative of active chromatin (Dixon et al. 2012; Ulianov et al. 2016) and genes within TADs are co-transcriptionally regulated (Nora et al. 2012; Galupa and Heard 2017; Li et al. 2018). TADs were implicated in replication activation mechanisms mediated by the spatial organization of the surrounding DNA sequences (Pope et al. 2014; Sima et al. 2019). However, if and how TADs affect the positioning and firing of replication origins is still unknown.

In this work, we explore the localization of replication origins firing in early S-phase and identified by EdU-seq, relative to TADs. In addition to mapping CN origins, EdU-seq technology has shown that oncogene overexpression triggers illegitimate origins which fire within highly transcribed genes, ultimately leading to fork collapse and DNA damage accumulation (Macheret and Halazonetis 2018; Macheret et al. 2020). Here we took advantage of the high-resolution origin mapping by EdU-seq to demonstrate that early S-phase CN origins significantly correlate with TAD boundaries. The latter were mapped using data obtained in 25 different cell lines. In contrast, oncogene-induced (Oi) origins were depleted in boundary regions. Moreover, we show that chemical transcription inhibitors deregulate the observed origin firing sites relative to TADs, confirming that transcription plays a key role in determining the sites of early origin firing. Following release from hydroxyurea (HU) block replication re-starts bi-directionally from forks within TAD boundaries. We bring evidence that forks move towards the centre of the TAD unobstructed. However, forks moving outwards from the TAD boundary are significantly constrained by the proximity of divergently-oriented CTCF binding sites or of TADs with different RT. Therefore, forks initiating at CN origins within TAD boundaries show asymmetrical progression after restart from HU arrest.

## RESULTS

### Qualitative differences in the mapping and timing of CN and Oi replication origins

The EdU-seq assay has previously been used to monitor DNA replication initiation in U2OS human osteosarcoma cells (Macheret and Halazonetis 2018). It relies on arresting cells with nocodazole in mitosis, followed by mitotic shake-off and releasing cells synchronously in the cell cycle in the presence of EdU and HU. Addition of HU limits the analysis to origins firing in early S-phase. Then, isolated EdU-labelled DNA is subjected to high-throughput sequencing allowing to identify the genomic regions of EdU incorporation (Macheret and Halazonetis 2018).

Here, we retrieved the sequencing data from the afore-mentioned study and re-analysed it in order to obtain genome-wide maps of CN origin firing sites and aberrant firing events induced by over-expressing cyclin E (*CCNE1*) in U2OS cells (Macheret and Halazonetis, 2018). The EdU-seq signal displayed well-resolved (10 Kb resolution) peaks which correspond to CN or Oi origin firing sites (top and middle panels, respectively; **Figure 1A,B**). CN origins (*n* = 3,956) were defined as origins that fired with similar intensity in U2OS cells expressing normal or high levels of cyclin E, while sites displaying peaks 4-fold higher when cyclin E was over-expressed compared to normal cells were defined as Oi origins (*n* = 927) (**Figure 1B**). A similar analysis was performed in human non-small cell lung carcinoma H1299 cells with either normal or induced expression of the β-catenin oncogene (**Figure S1** and unpublished data).

**Figure 1.**
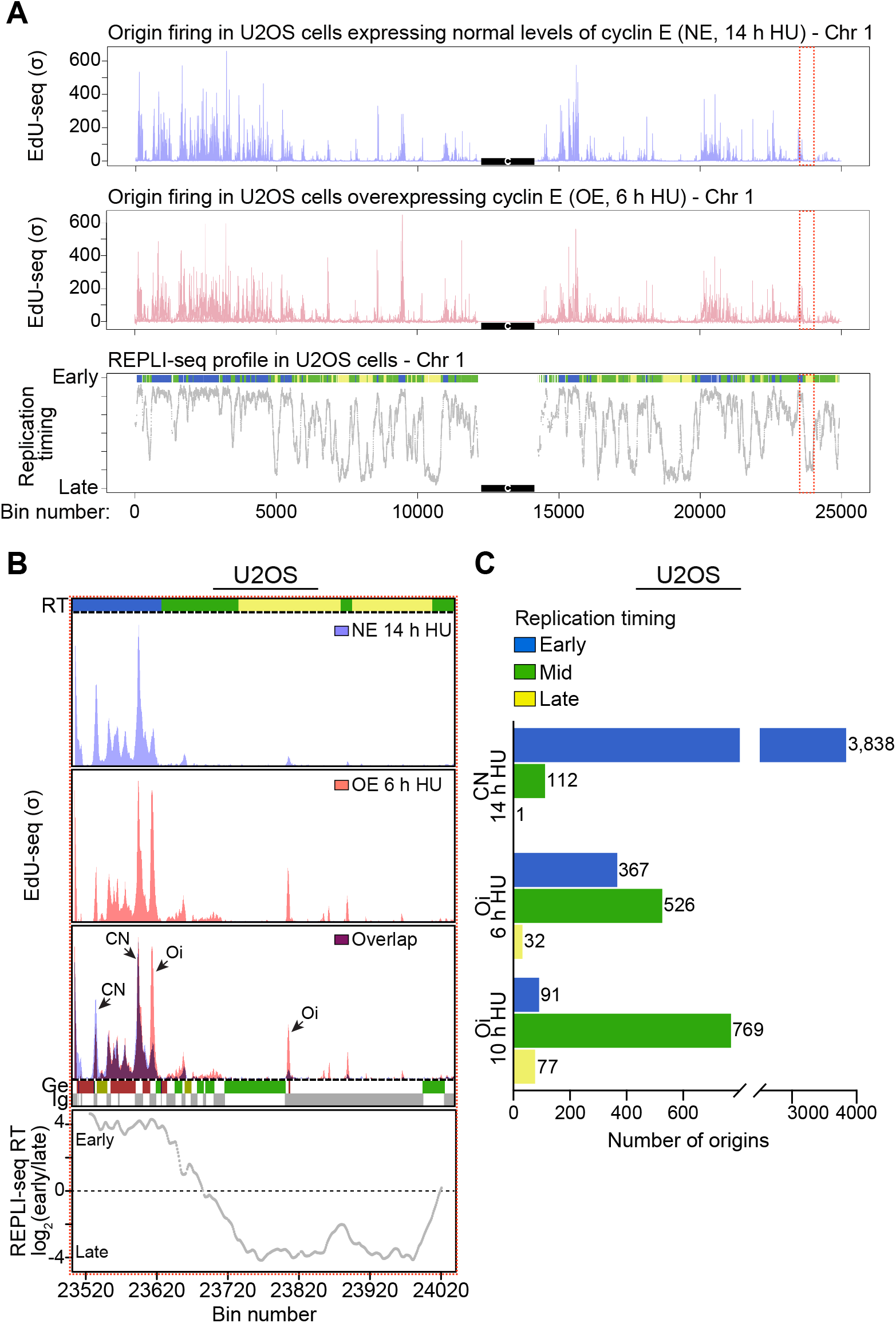
Quantification of localization and timing of replication origin firing in U2OS cells. **(A)** Distribution of origin firing events along chromosome 1 based on EdU-seq signal (σ, normalized number of sequence reads per 10 Kb bin divided by its standard deviation) in U2OS cells expressing normal levels of cyclin E (NE), collected 14 h after mitotic shake-off (light purple) or overexpressing cyclin E (OE), collected 6 h after mitotic shake-off (red). Distribution of RT domains detected by REPLI-seq along chromosome 1 (grey; Loess-smoothed RT profiles per 10 Kb-bins). Top bar indicates S-phase timing: blue, early; green, mid; yellow, late. Raw data retrieved from (Macheret and Halazonetis 2018). Black boxes indicate the centromere position on chromosome 1. **(B)** High-resolution representation of the region indicated by the red box in **(A)**. RT, replication timing (blue, early; green, mid; yellow, late S-phase); CN, constitutive origin; Oi, oncogene-induced origin; Ge, genic (green, forward direction of transcription; red, reverse; yellow, unspecified; blue, multiple genes within bin); iG, intergenic (grey). Bin resolution, 10 Kb. **(C)** Genome-wide quantification of CN (14 h after mitotic shake-off) and Oi (6 or 10 h after mitotic shake-off) according to their RT.

Next, we compared the genome-wide distribution of CN and Oi origins relative to RT domains identified with REPLI-seq (Hansen et al. 2010). We retrieved REPLI-seq data obtained in U2OS cells (Macheret and Halazonetis 2018), and mapped early, mid or late S-phase replicating domains (bottom panel; **Figure 1A**). Previous studies have shown that timing is largely conserved among cell types, resulting in highly reproducible RT domains (Hiratani et al. 2008; Hansen et al. 2010; Ryba et al. 2011; Fragkos et al. 2015). Thus, the assignment of RT domains used for reference in all our analyses was based on REPLI-seq profiles obtained in U2OS cells. CN origins mapped by EdU-seq were present almost exclusively in early replicating domains (97% of all identified origins; **Figure 1C**). In contrast, Oi origins showed a wider distribution, as they fired both within early (40% of Oi origins, 6 h after mitotic shake-off) and mid S-phase (57% of Oi origins, 6 h after mitotic shake-off) domains. Origin quantification revealed that a fraction of the Oi origins fired in mid S-phase and this increased with S-phase progression (*n* = 526 mid S Oi origins or 57% of all identified origins at 6 h after mitotic shake-off; *n* = 790 mid S Oi origins at 10 h after mitotic shake-off or 82% of all identified origins; **Figure 1C**).

Analysis of RT revealed differences in origin firing timing between U2OS and H1299 cells. A higher fraction of H1299 CN mapped to mid S-phase domains, 12 h after release from mitotic arrest (*n* = 867 or 29% of all detected CN origins; **Figure S1B**). This could be attributed to a shorter G1 phase in H1299 cells compared to U2OS cells. Similarly to U2OS cells overexpressing cyclin E, the relative distribution of Oi origins induced by β-catenin overexpression in H1299 cells was shifted towards mid S-phase replication (*n* = 186 mid S-phase Oi origins at 12 h after mitotic shake-off or 63% of all identified origins; **Figure S1B**).

### CN and Oi origins differ in their genomic distribution relative to TADs

CTRs coincide with TAD boundaries (**Figure S2A**), thus these discrete genomic entities may play a role in defining RT (Moindrot et al. 2012; Pope et al. 2014; Marchal et al. 2019). Given the observed temporal differences between CN and Oi origin firing, we sought to compare their positions relative to TADs.

We first defined TADs based on maps retrieved from the 3D Genome Browser for 25 different human cell lines (**Figure S2B**). Importantly, all TAD maps were systematically predicted from Hi-C data using the same hidden Markov model (HMM) pipeline (Dixon et al. 2012). This enabled us to avoid variation in TAD sizes and numbers due to previously reported discrepancies in computational identification methods (Dali and Blanchette 2017; Forcato et al. 2017; Zufferey et al. 2018). The median TAD length across the 25 cell lines analysed was 920 Kb (**Figure S2C**) and the median inter-TAD distance (defined as the distance between the end coordinate of a given TAD and the start coordinate of the subsequent one) was 40 Kb (**Figure S2D**). We therefore defined TAD boundaries as the 40 Kb window surrounding (± 20 Kb) the start and end coordinates of each TAD. This window size is consistent with TAD boundaries defined by other studies (Gong et al. 2018; McArthur and Capra 2020), the calculated median inter-TAD distance and the Hi-C resolution of TAD maps (25 to 40 Kb range), thus accounting for the uncertainty in the exact start/end of TAD coordinates (Wang et al. 2018; Zufferey et al. 2018).

We then mapped genome-wide the average EdU-seq signal (σ) for CN and Oi origins firing in U2OS cells, relative to TADs defined as above. We observed that the signal intensity corresponding to CN origins firing in early S-phase peaks within TAD boundaries gradually decreases towards the centre of TADs (**Figure 2A**). In striking contrast to this pattern, Oi origins fired predominantly in the centre of TADs, with only residual EdU-seq signal within boundaries (**Figure 2B**). Transcription before S-phase entry has been proposed to inactivate aberrant origin firing within genes (Sasaki et al. 2006; Powell et al. 2015). Consistent with this, treatment with 5,6-dichloro-1-β-d-ribofuranosylbenzimidazole (DRB), a chemical inhibitor of transcription elongation, of U2OS cells in G1 has been shown to trigger aberrant origin firing at the same loci as Oi origins (Macheret and Halazonetis 2018). Thus, we examined origin firing distribution in U2OS cells expressing normal levels of cyclin E, treated with DRB for the first 9 h of release from mitotic arrest, when cells are in G1. Our analyses showed that DRB-induced origins (DRBi) had the same distribution as Oi origins relative to TADs, being enriched in the centre of TADs and depleted in TAD boundaries (**Figure 2C**).

**Figure 2.**
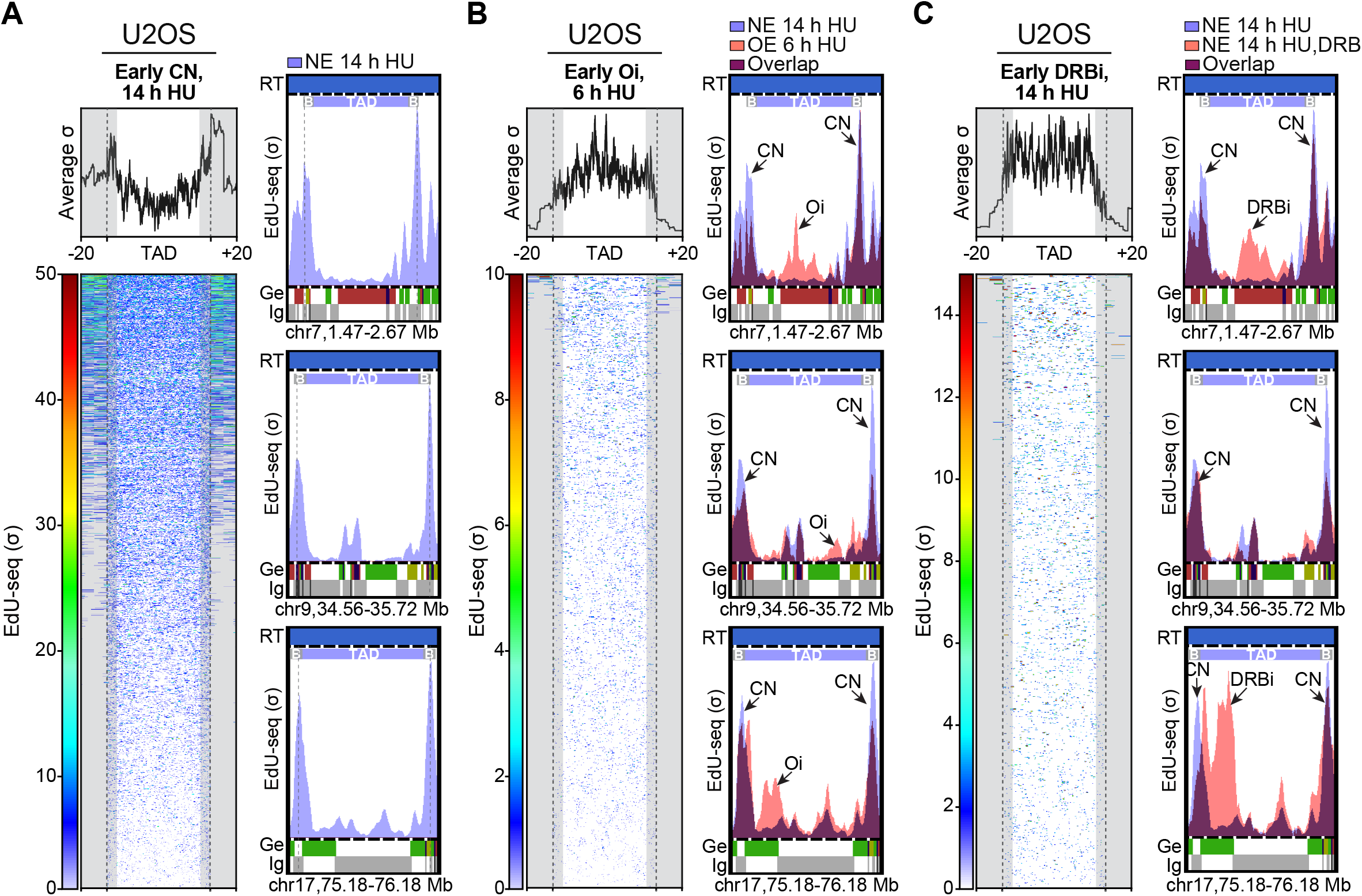
Distribution of replication origins firing in early S-phase relative to TADs in U2OS cells. **(A-C)** Genome-wide early replication origin firing profiles (based on average EdU-seq σ values) are represented relative to TAD positions, with TAD boundaries (±20 Kb from TAD start/end points indicated by dotted lines) shadowed in grey. Heatmaps represent EdU-seq σ values over each of the inspected TADs. Representative windows of replication initiation profiles on chromosome 7, 9 and 17 are shown for each average σ graph. **(A)** Early constitutive (CN) origins were mapped relative to the average position of *n* = 21,226 TADs having at least 1 early origin. **(B)** Early oncogene-induced (Oi) origins were mapped relative to the average position of *n* = 6,295 TADs having at least 1 early origin. **(C)** Early origins induced by DRB treatment (DRBi) of G1 cells (0-9 h after release form mitotic arrest) were mapped relative to the average position of *n* = 4,716 TADs having at least 1 early origin. B, TAD boundary; RT, replication timing (blue, early S-phase); σ, normalized number of sequence reads per 10 Kb-bin divided by its standard deviation; Ge, genic (green, forward direction of transcription; red, reverse; yellow, unspecified; blue, multiple genes within bin); iG, intergenic (grey). TAD and TAD boundary mapping was based on average data obtained from 25 human cell lines shown in **Figure S2**.

TAD boundaries are enriched in housekeeping genes (Ciabrelli and Cavalli 2015) and histone marks characteristic of actively transcribed chromatin (Dixon et al. 2012; Ulianov et al. 2016). Consistent with this, we found that in our U2OS model there is a significant overlap of housekeeping genes with TAD boundary regions (1,212 genes out of the 9,786 genes located entirely within or partially overlapping TAD boundaries, Fisher’s exact test *P* = 3.4×10^−21^; see *Methods* for the definition of the housekeeping gene set). However, it is not clear whether the genes located within TAD boundaries are actively transcribed at the time of early CN origin firing. To address this, we compared the profiles of 5-ethynyl-uridine (EU) incorporation genome-wide using the average EU-seq signal for newly synthesized transcripts relative to TADs in U2OS cells, using the same timing as for EdU-seq. We observed that the highest EU-seq signal mapped to TAD boundaries, regardless of the direction of transcription and in both normal U2OS cells or those overexpressing cyclin E (**Figure S3A,B**). Oi origins fire in regions displaying the lowest transcription levels within TADs (**Figure S3B**). Although CN origins preferentially fire within TAD boundaries, we observed distinct peaks of the EdU-seq and EU-seq signal within these regions, suggesting that the sites of replication initiation do not coincide with those of active transcription (**Figure S3A,C**). It is possible, however, that the proximity of active transcription promotes origin firing, by remodelling the chromatin into a configuration that allows ORC binding. Moreover, we found that approximately 80% of the TAD boundaries (**Figure S3D**) are located within intergenic regions. This is also consistent with the previous observation that early CN origins map predominantly to intergenic regions, thus minimizing possible replication–transcription conflicts (Martin et al. 2011; Macheret and Halazonetis 2018).

### Early CN origins are specifically enriched within and adjacent to TAD boundaries

Our analyses showed that CN and Oi origins are differently positioned within TADs, with a remarkable enrichment of early CN origins within the TAD boundary. Boundaries are insulator elements that contribute to the restriction of promoter–enhancer communication to each TAD (Shen et al. 2012; Symmons et al. 2014) and are enriched in cohesin complex and CTCF binding sites (Dixon et al. 2012; de Wit et al. 2015; Nora et al. 2017; Rao et al. 2017).

We further strengthened this analysis by calculating the number of detected origins within a 800 Kb window surrounding (±400 Kb) the TAD boundary. Genome-wide boundaries (start and end) were assessed together. Our results indicated that early CN origins fire most frequently within or adjacent to TAD boundaries genome-wide (**Figure 3A**; permutations test for association evaluation, *P* < 1×10^−4^, *n* = 10,000 circular permutations). In contrast with this, neither Oi or DRBi origins were significantly associated with TAD boundaries, being most frequently positioned outside these regions (**Figure 3A**; permutations test for association evaluation, *P* >0.05, *n* = 10,000 circular permutations). Next, we increased the resolution of our analysis by representing the frequency of early S-phase origin firing around TAD boundaries using smaller genomic windows (±120 Kb around TAD boundaries). Interestingly, for CN origins we resolved two distinct peaks mapping to the edge or in the proximity of the TAD boundary (**Figure 3B**). We hypothesized that the position of CN firing peaks could be due to structural TAD components binding to the centre of the boundary and thus preventing origin activation. We tested this hypothesis by examining ChIP-seq data for the transcription factor CTCF, and for cohesin components RAD21 and SMC3. Consistent with previous reports, we found that TAD boundaries are significantly enriched in binding sites for these factors. We detected maximal CTCF, RAD21 and SMC3 binding peaks at the centre of TAD boundaries (**Figure 3C**; permutations test for each association, *P* < 1×10^−4^, *n* = 10,000 circular permutations), flanked by EdU-seq peaks. One possible explanation for this positioning is that active transcription emanating from CTCF binding sites displaced replication initiation sites towards the edges of the TAD boundary.

**Figure 3.**
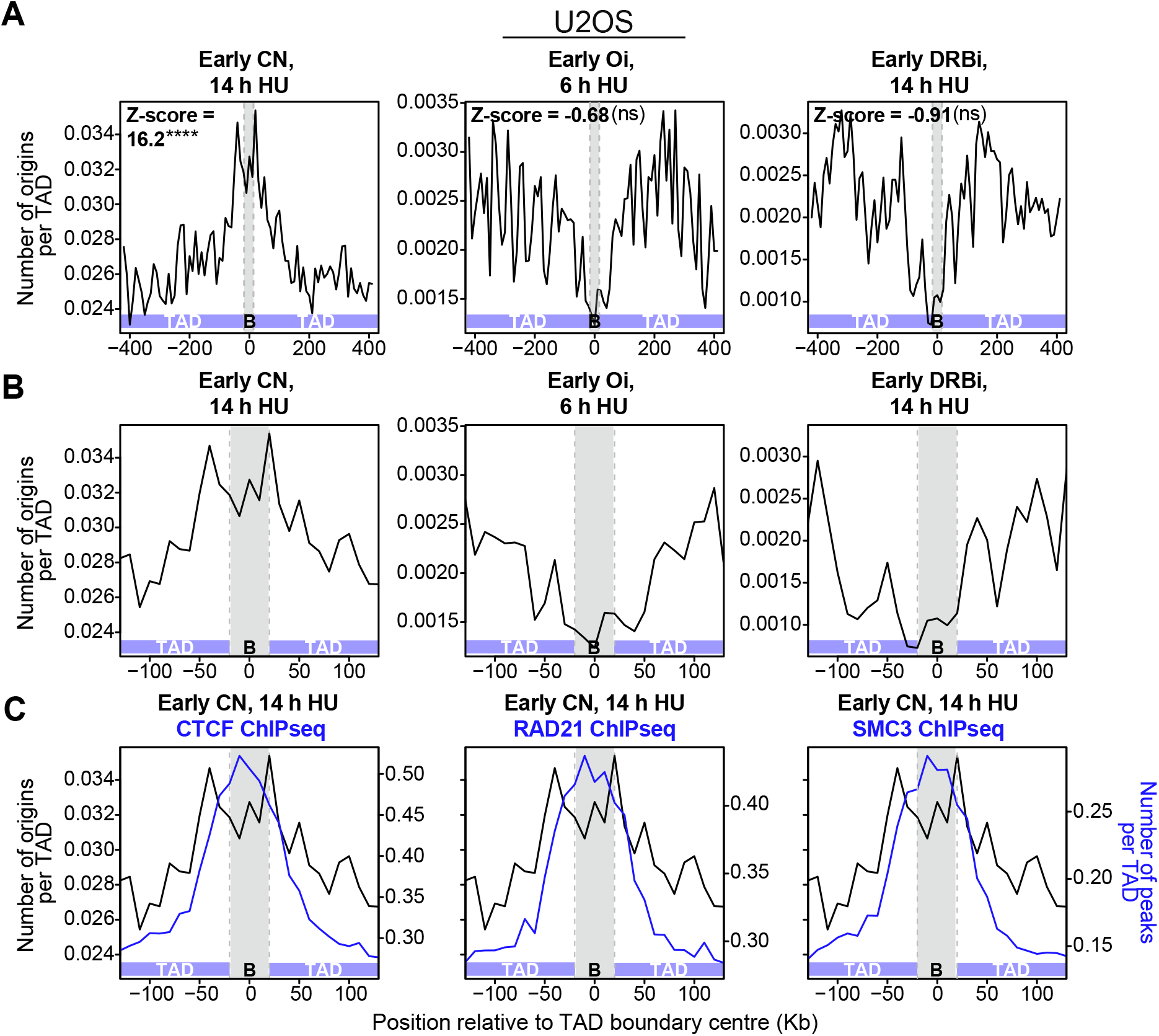
Mapping early replication origins relative to TAD boundaries in U2OS cells. **(A)** Frequency of early replication origins firing within TAD boundaries and adjacent regions of ±400 Kb. Positive z-scores indicate that origins are enriched in TAD boundaries, whilst negative z-scores indicate that origins are depleted from TAD boundaries. ****, *P* < 1×10^−4^; ns, *P* > 0.05, permutations tests for association (*n* = 10,000 circular permutations). CN, constitutive origins; Oi, oncogene-induced origins; DRBi, origins induced by DRB treatment of G1 cells (0-9 h after release form mitotic arrest). B, TAD boundary. **(B)** Frequency of early replication origins firing within TAD boundaries and adjacent regions of ±120 Kb. Origins were classified as in **(A)**. **(C)** Frequency of CN early replication origins (black) and ChIP-seq peaks (blue) within TAD boundaries and adjacent regions of ±120 Kb. ChIP-seq signal, from left to right: CTCF ChIP-seq signal was calculated based on overlapping data available from 8 cell lines (z-score = 40.9; *P* < 1×10^−4^, permutations tests for association using *n* = 10,000 circular permutations); RAD21 ChIP-seq signal was calculated based on overlapping data available from 6 cell lines (z-score = 29.1; *P* < 1×10^−4^, permutations tests for association using *n* = 10,000 circular permutations). SMC3 ChIP-seq signal was calculated based on overlapping data available from 4 cell lines (z-score = 50.2; *P* < 1×10^−4^, permutations tests for association using *n* = 10,000 circular permutations). The details of ChIP-seq datasets used are shown in **Table S1**.

The results obtained in U2OS were recapitulated in H1299 cells, where analyses of average EdU-seq signal and origin frequency indicated that early CN origins were located within TAD boundaries, whilst Oi (β-catenin-induced) origins were located in the middle of the TAD (**Figure S4A,C**). H1299 cells progress faster into S-phase, which enabled us to capture some origin firing events in mid S-phase. CN mid S-phase origins were enriched at TAD boundaries, whilst Oi mid S-phase origins fired more frequently towards TAD centres (**Figure S4B,D**), similarly to the early origins. Origin frequency peaks were observed at TAD boundary edge or in its vicinity, for both early and mid S-phase CN origins in H1299 cells (**Figure S4C,D**). Altogether, our analyses indicate that CN and Oi origin firing events mapped by EdU-seq differ not only in their timing, but also in their positioning in the three-dimensional genome space. Their genomic position relative to TAD boundaries is conserved between different human cell lines and overexpressed oncogenes.

Next, we sought to determine the positions of early CN origins relative to TADs using a different sequencing technology for origin mapping. We focused on SNS-based methods, which rely on purification of nascent DNA strands specific to replication origins (1.5 to 2.5 Kb in size) based on their resistance to λ-exonuclease digestion (Bielinsky and Gerbi 1998). Using high-throughput sequencing in asynchronous HeLa cells, the range of replication origins detected genome-wide has been expanded to approximatively 250,000 origins (Besnard et al. 2012). Moreover, previous studies detected a significant overlap between cohesin binding sites and replication origins identified by SNS-seq (Cadoret et al. 2008; Guillou et al. 2010). Previous studies have reported that purification of short nascent strands using the λ-exonuclease assay could intrinsically bias origin enrichment towards GC-rich DNA, notably G-quadruplexes, which cannot be efficiently digested by this enzyme (Foulk et al. 2015; Petryk et al. 2016). Thus, we reanalysed the data from Besnard *et al.* using non-replicating genomic DNA (LexoG0 data) as a control (Foulk et al. 2015). This enabled identification of a total of 78,731 origins in HeLa cells (**Figure S5A,B**). Consistent with the origin maps determined by EdU-seq, both early and mid S-phase SNS origins were significantly enriched at TAD boundaries (**Figure S5C**; permutations tests for association evaluation, *P* < 1×10^−3^ and *P* < 1×10^−4^, respectively, *n* = 10,000 circular permutations). However, we did not observe an enrichment of late SNS origins at these sites (negative z-score, permutations test for association evaluation, *P* > 0.05, *n* = 10,000 circular permutations; data not shown), possibly due to the small number of detectable origins in this group.

### Asymmetrical fork progression at TAD boundaries after HU release

Having established that early CN origins fire more frequently within or adjacent to TAD boundaries, we investigated next whether this localization could impact on the restart and progression of replication forks after release from HU arrest. EdU-seq release experiments rely on restarting arrested forks by removing HU from the media and monitoring fork progression using EdU incorporation during release. We thus analysed EdU-seq release data obtained in U2OS cells (Macheret and Halazonetis 2018) for early CN origins firing within TAD start/end boundaries or within the TAD itself (**Figure 4A**). Forks originating inside TADs recovered and advanced symmetrically both at 90 min and 150 min after HU release (**Figure 4A**, middle panel). In striking contrast, forks originating at TAD boundaries showed a clear asymmetry as early as 90 min after release from HU block, which became more pronounced 150 min after release (**Figure 4A**, left and right panels). Specifically, forks moving outwards from either the start or end TAD boundaries showed slower progression, indicative of fork stalling. The same asymmetry in fork progression from early CN origins within TAD boundaries as in U2OS was observed in H1299 cells treated with HU for 12 h and released for 120 min, as illustrated in representative images of forks firing at a selected region on chromosome 1 (**Figure 4B**). Conversely, progression of forks originating at CN origins within the TAD was unperturbed and symmetrical in both U2OS and H1299 cells (**Figure 4A,C**). Consistent with a previous report (Macheret and Halazonetis 2018), replication forks initiating at Oi origins located in TAD centre failed to progress and collapsed shortly after firing (**Figure S6A,B**).

**Figure 4.**
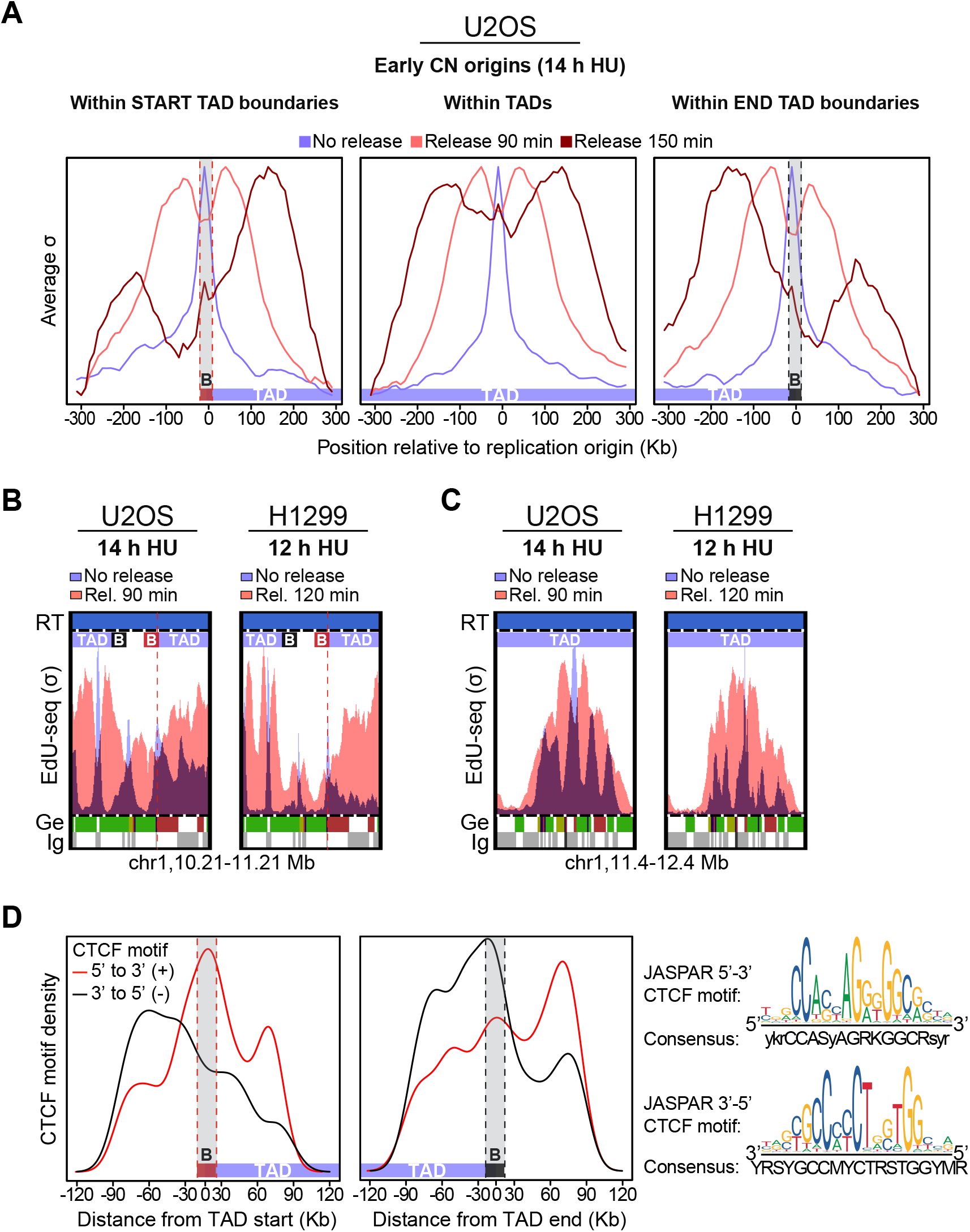
Restart and progression of replication forks originating in TAD boundaries after release from HU arrest. **(A)** Average EdU-seq σ values showing early origin firing in U2OS cells treated with HU for 14 h (light purple) and fork progression in U2OS cells released from HU arrest for 90 min (light red) or 150 min (dark red). Early CN origins were mapped to start TAD boundaries (*n* = 603) non-overlapping with a preceding end boundary (left), TADs (*n* = 1,242) non-overlapping with either start or end boundaries (middle) and end TAD boundaries (*n* = 535) non-overlapping with a subsequent start boundary (right). B, TAD boundary. **(B)** Representative genomic region (chr1:10,210,000-11,210,000 bp) showing a CN origin firing within TAD boundary. U2OS cells were treated with HU for 14 h, followed by release for 90 min. H1299 cells were treated with HU for 12 h, followed by release for 120 min. RT, replication timing (blue, early S-phase); σ, normalized number of sequence reads per 10 Kb-bin divided by its standard deviation; Ge, genic (green, forward direction of transcription; red, reverse; yellow, unspecified; blue, multiple genes within bin); iG, intergenic (grey); bin resolution, 10 Kb. (**C**) Representative genomic region (chr1:11,400,000-12,400,000 bp) showing a CN origin firing within TAD. EdU-seq σ values indicate origin firing (light blue) and asymmetric fork progression (light red) in U2OS and H1299 cells treated as in **(B)**. **(D)** Density of the most robust CTCF motifs (FDR < 0.05, 3rd quartile) mapping within ±120 Kb from start (left) or end (right) TAD boundaries. Densities were calculated independently for CTCF motifs oriented 5’-to-3’ (red) and 3’-to-5’ (black). The consensus sequence used for the search of CTCF sites is represented in the bottom (JASPAR core CTCF motif, MA0139.1). B, TAD boundary.

### Barriers to fork progression in the vicinity of TAD boundaries

TADs are postulated to form by an active loop extrusion mechanism, which requires binding of the CTCF protein and the cohesin complex at both boundaries (Sanborn et al. 2015; Fudenberg et al. 2016; Rao et al. 2017). The cohesin complex associates randomly with chromosomal DNA and promotes loop extrusion in bidirectional manner, until an inward, convergently-oriented CTCF-bound site is encountered. Loop extrusion is completed when both sides of the loop encountered convergent CTCF binding sites (Fudenberg et al. 2016; Davidson et al. 2019; Kim et al. 2019). We hypothesized that the asymmetric progression of forks initiating from origins located within TAD boundaries could be due to CTCF binding that restrict the outwards progression, causing fork stalling. We determined computationally the densities of CTCF binding sites (JASPAR core database, MA0139.1) within a 240 Kb window surrounding (±120 Kb) start/end TAD boundaries (**Figure 4D**). Our results showed that 5’-to-3’convergently-oriented sites peak within the start TAD boundaries and 3’-to-5’ convergently-oriented sites peak within the end TAD boundaries, as predicted by the loop extrusion model. However, the region preceding the start boundary was enriched in 3’-to-5’ divergently-oriented motifs, whilst the region beyond the end boundary was enriched in 5’-to-3’ divergently-oriented motifs. Because fork progression is obstructed specifically in the TAD external regions immediately adjacent to the boundary (**Figure 4A**), we propose that the divergently-oriented CTCF sites represent physical barriers to replication.

We observed that only a fraction of forks initiating from early CN origins within TAD boundaries progressed asymmetrically. For example, approximately 31% of non-overlapping 800 Kb windows on chromosome 1 displayed replication forks with asymmetric progression and 60% of these coincided with regions where REPLI-seq determined RT switched from early to mid S-phase (**Figure 5A**; red arrows). To test whether the timing transition regions could also obstruct replication, we analysed the EdU-seq release signal around TAD boundaries, whilst concomitantly defining the RT of adjacent TADs. Consistent with our previous data, the forks originating in either start or end boundaries progressed preferentially towards the centre of the TAD, but frequently stalled when entering the neighbouring TAD with a different RT (**Figure 5B,C**). Therefore, we propose that RT of adjacent TADs is a contributing factor, in addition to CTCF binding sites, to the asymmetry of forks initiating from early CN origins within TAD boundaries (Figure 6).

**Figure 5.**
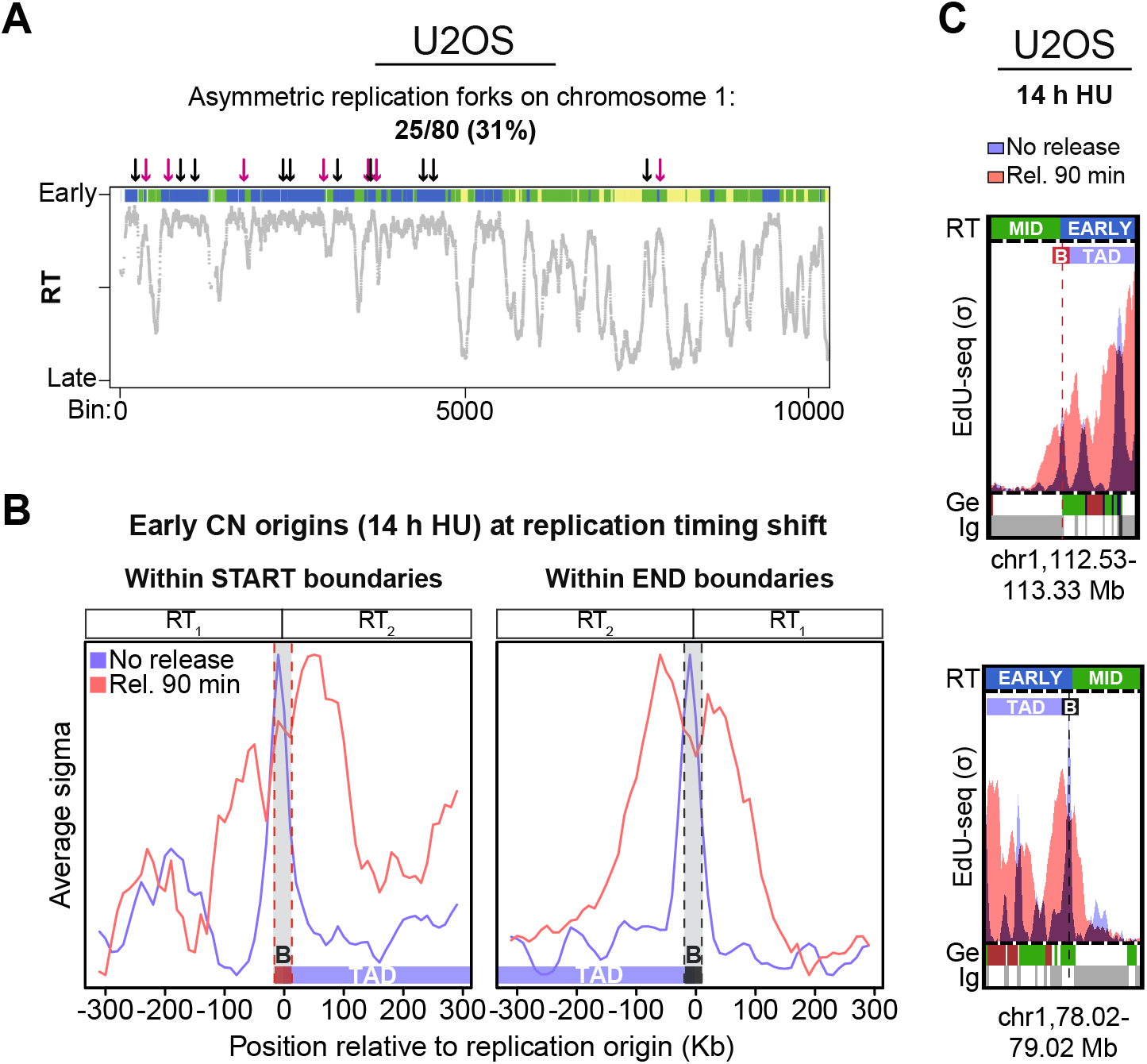
Replication fork restart and progression at RT transition regions. **(A)** Non-overlapping 80-bin (800 Kb) windows, containing at least one replication fork, on chromosome 1 (first 10,000 bins, p-arm) were inspected for asymmetries in the EdU-seq release signal around early S-phase CN origins firing in TAD boundaries. The arrows indicate the positions of the asymmetric forks detected (31% of the inspected windows) along chromosome 1; purple arrows indicate windows with asymmetric forks at RT shifts. **(B)** Average EdU-seq σ values showing early origin firing in U2OS cells treated with HU for 14 h (light blue) and fork progression in U2OS cells released from HU arrest for 90 min (light red). Early CN origins were mapped to: start TAD boundaries overlapping a RT shift (*n* = 124); or end TAD boundaries overlapping a RT shift (*n* = 139). B, TAD boundary; RT, replication timing. **(C)** Representative genomic regions showing CN origin firing and asymmetric fork progression around start (chr1:112,530,000-113,330,000 bp) and end TAD boundaries (chr1:78,020,000-79,020,000 bp). U2OS cells were treated with HU for 14 h, followed by release for 90 min. RT, replication timing (blue, early S-phase); σ, normalized number of sequence reads per 10 Kb-bin divided by its standard deviation; Ge, genic (green, forward direction of transcription; red, reverse; yellow, unspecified; blue, multiple genes within bin); iG, intergenic (grey); bin resolution, 10 Kb.

**Figure 6.**
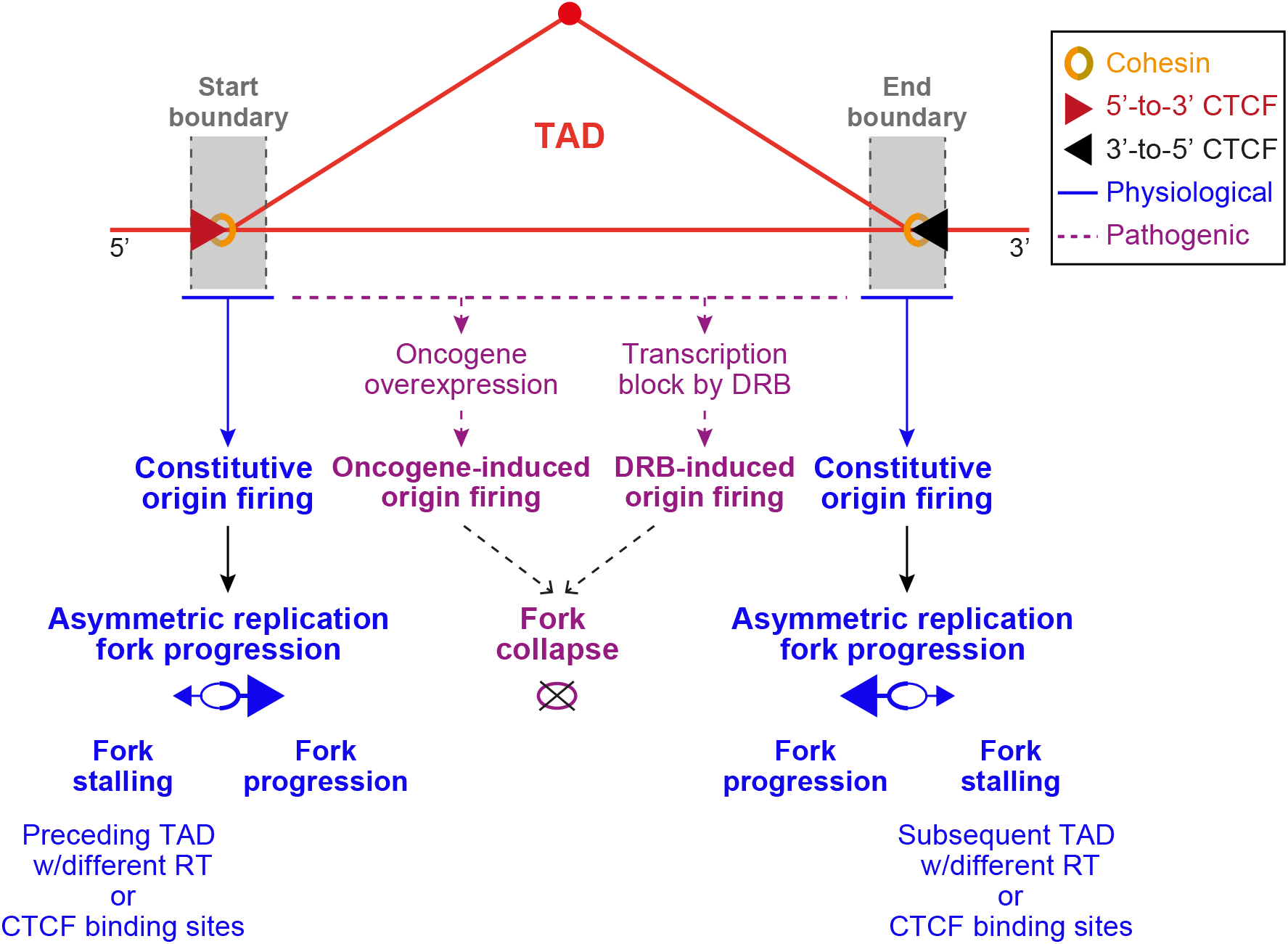
Model for CN and Oi origin firing and fork progression relative to TADs. Under physiological conditions, early S-phase CN replication origins fire with highest frequency in or in the vicinity of TAD boundaries. Boundary regions are enriched for both convergent CTCF (5’-to-3’ inwards oriented motif at the start of the TAD and 3’-to-5’ oriented at its end) and cohesin complex binding sites. In addition, a fraction of TAD boundaries also colocalize with RT transition regions (from/to early S-phase). These features impact on the progression of replication forks initiating at early S-phase origins, which results in asymmetric fork movement preferentially towards the centre of the TAD. Outward movement is blocked by CTCF sites and/or the vicinity of a TAD with different RT. Pathogenic conditions, such as oncogene overexpression or DRB-dependent transcription block, trigger origin firing predominantly within the TAD and lead to fork collapse shortly after firing.

## DISCUSSION

In human cells, replication origin activation had initially been predicted based on ORC binding, where the replicative helicase MCM2–7 is loaded onto DNA (Bell and Kaguni 2013). ORC binding sites are frequently positioned within euchromatin regions characterised by DNase I hypersensitivity (Gindin et al. 2014; Cayrou et al. 2015; Miotto et al. 2016) and therefore lack sequence specificity. ORC loading relies on opportunistic binding to DNA sites where chromatin was made accessible by active transcription. Additionally, the chromatin context, in particular histone modifications around replication origins, is important in determining RT during S-phase. Indeed, the probability of a specific origin firing can be enhanced or suppressed depending on which histone acetylases or deacetylases are recruited in its vicinity (Vogelauer et al. 2002; Goren et al. 2008; Mantiero et al. 2011). This suggested that the timing of origin firing is not pre-set, but its local chromatin context can promote or suppress its activation during early, mid or late S-phase.

A consensus map of replication origin positions has not yet been assembled, as there are major inconsistencies between maps obtained using different high-throughput methods. Consequently, the number of putative origins in the human genome is ranging from 4,000 (Macheret and Halazonetis 2018) to around 250,000 (Besnard et al. 2012). Moreover, the genomic positions of the active origins differ between these approaches: for instance, 33 to 65% overlap was found between SNS-seq and Bubble-seq origins, depending on the peak calling parameters used (Hyrien 2015; Langley et al. 2016). To date, the precise factors that specify which origins fire in any given S-phase remain poorly understood (Prioleau and MacAlpine 2016) and no universal signature has yet been identified to fully predict the origins active at a given time in mammalian genomes.

Recent computational models based on high-throughput approaches carried out in mammalian cells have characterized large genomic regions with relatively uniform RT, or CTRs (Farkash-Amar et al. 2008; Ryba et al. 2010; Guilbaud et al. 2011; Rhind and Gilbert 2013)). These chromosomal domains are characterized by synchronous origin firing within clusters of replication initiation events (Hiratani et al. 2008; Pope et al. 2014; Marchal et al. 2019). In particular, replication domain borders, defined as sites where RT transitions from early to late, have recently gained substantial interest because they frequently align with TAD boundaries (Pope et al. 2014). Further supporting the link between the RT programme and the three-dimensional organization of the genome into TADs was the finding that both are established in early G1 (Dimitrova and Gilbert 1999; Dileep et al. 2015). Thus, there is a strong correlation between genome architecture and RT of a given chromosome domain, but no causal link has been directly established and no direct connection between origin activity and TAD boundary elements has been found.

Here we used high-resolution (10 Kb) genome-wide maps of early replication origins generated by EdU-seq in two different human cell lines (U2OS and H1299), to determine their position relative to TAD boundaries. The robustness of TAD positions, which are not as stable as initially thought (Fudenberg and Pollard 2019; McArthur and Capra 2020; Sauerwald et al. 2020), was ensured by averaging a collection of TAD maps obtained from 25 cell lines. Our results demonstrate that genome-wide EdU-seq origin firing profiles display enrichment of early CN origins within or in the proximity of 40 Kb TAD boundary regions. We demonstrated the preferential localisation of early CN origins within TAD boundaries also using SNS-seq origin mapping in HeLa cells.

In striking contrast with the position of early CN origins, we find that early Oi origins fire within the TAD. Transcription inhibition in G1 using DRB triggered the same origin localisation pattern as Oi. Both DRBi and Oi origins fire within genes (Macheret and Halazonetis 2018), whilst CN origins fire preferentially in intergenic regions. Although S-phase transcription profiles determined with EU-seq also peak in TAD boundaries, these EU-seq peaks do not coincide with the EdU-seq peaks for early CN origin firing. Therefore, replication and transcription initiate at distinct sites within the TAD boundary. Moreover, we find that G1 transcription is required to specifically activate early origins within the boundary. If G1 transcription is abrogated, alternative origins are activated, which are positioned within the TAD and within actively transcribed genes. Taken together, these results suggest that transcription within TAD boundaries is a pre-requisite for early origin firing. Conceivably, transcription-associated chromatin remodelling within the boundary could also promote replisome assembly.

The EdU-seq technology enabled us to monitor the restart after HU arrest and progression of replication forks initiating at CN origins localised within TAD boundaries. Whilst normal fork movement is bidirectional, a subset of forks originating at TAD boundaries showed a striking asymmetry, with preferential progression towards the centre of the TAD. We identified two mechanisms that obstruct fork progression outside TAD boundaries: divergent CTCF binding sites and the proximity of a TAD with different RT (**Figure 6**). Consistent with this, we did not detect any asymmetry in forks arising from the subset of CN origins that fire within TADs, where they would not encounter either of these physical barriers. Oi origins fire within the TADs at high frequency, but they fail to progress and collapse shortly after restart. This could be due to their propensity to localise within actively transcribed genes, whilst CN origins largely localise in intergenic regions.

Our results support the concept that TADs represent not only genome architecture units, but also RT entities. Recent work (Sima et al. 2019) proposed that CTCF-dependent genomic architecture is dispensable for the execution of RT patterns at a specific genomic locus, thus arguing against the hypothesis that TADs act as RT organization subunits. Our results are consistent with this because we find that both early and mid S-phase origins fire within TAD boundaries in HeLa and H1299 cells. Therefore, TAD boundaries do not necessarily define the timing of origin firing. However, they provide a chromatin environment conducive for the recruitment of the replication machinery, which requires CTCF-dependent initiation of transcription and chromatin remodelling.

It remains unknown why fork progression is impaired at the transition between TADs with different RT and whether the enrichment in CTCF binding sites divergently orientated external to the TAD is a contributing factor (Figure 6). Nevertheless, these features affect a significant subset of the replication forks within the genome. Understanding their aetiology could provide insight into why some parts of the genome are more difficult to replicate than others. Replication failure at these sites could, in turn, lead to chromosome rearrangements that drive genomic instability. Therefore, our work demonstrates for the first time that the three-dimensional genomic architecture could act as a natural barrier to replication fork progression and pose a threat to chromosome integrity.

## METHODS

### EdU-seq and EU-seq data processing

Raw FASTQ files were retrieved from the Sequence Read Archive (SRA), accessions SRP126407 for U2OS cells (Macheret and Halazonetis 2018) and SRP269803 for H1299 cells (EdU-seq data only; unpublished). Sequence reads were aligned to the masked human genome assembly (GRCh37/hg19) using BWA-mem (Li and Durbin 2009), retaining only the reads with base quality scores ≥ 37. Then, the data was analysed as previously described (Macheret and Halazonetis 2019). Briefly, each chromosome was split into 10-Kb bins and the number of reads for each bin was determined. Next, sigma (σ) values for each genomic bin were calculated, as the normalized number of sequence reads per bin, divided by its standard deviation. For EU-seq data, samples were split by forward or reverse direction of transcription prior to σ calculation for each bin. For EdU-seq data, the obtained σ values were used to identify replication origins, by searching for local maxima in each chromosome. For a given origin, if the ratio of the oncogene (cyclin E or β-catenin)-overexpressed/control sigma values was greater than 4, then the origin was considered as oncogene-induced 4-fold (Oi, 4:1); similarly, if the ratio was greater than 2, but lower than 4, the origin was considered as oncogene-induced 2-fold (Oi2, 2:1). Unless otherwise stated, Oi 4-fold origins only were considered for all analyses. All other origins were considered to be constitutive (CN). Similarly, for DRB-induced origin firing (DRBi), if the ratio of the DRB-treated/no DRB control sample sigma values was greater than 4, then the origin was considered as induced by inhibition of transcription.

### Assignment of replication timing domains from REPLI-seq data in U2OS cells

Raw FASTQ files were retrieved from SRA, accession SRP126407 (Macheret and Halazonetis 2018). The early, mid and late S-phase REPLI-seq reads were assigned to 10-Kb genomic bins and corrected for variations in sequencing quality, as described in Macheret and Halazonetis 2018. Then, the numbers of early, mid and late S-phase reads were compared for each genomic bin: if one fraction (early, mid or late) accounted for more than 50% of the total reads for the bin, then that bin was assigned to the corresponding replication timing (RT) domain. The assignment of RT domains used for reference in all analyses was based on the U2OS samples expressing normal levels of cyclin E, which showed the sharpest REPLI-seq profiles. Finally, broad early/late RT domains were assigned following the protocol described by Marchal and colleagues (Marchal et al. 2018). Briefly, early (E) over late (L) RT was calculated as the log2(E/L) reads (within the 10-Kb bins) for the U2OS samples expressing normal levels of cyclin E. Duplicated reads were excluded from the analysis as well as reads mapped on chromosome Y or mtDNA. Quantile normalization and Loess smoothing with a span of 300 Kb per length of a given chromosome were applied to E/L RT values prior to generating the plots shown in **Figure 1**.

### SNS-seq replication origin maps in HeLa cells

Data for genome-wide sequencing of Short Nascent Strands (SNS) at replication origins in HeLa cells was retrieved from previously published work (Besnard et al. 2012). Raw FASTQ files were downloaded from SRA, accession SRX145994 (two replicates) and mapped to hg19 using BWA (Li and Durbin 2009). To control for reported nascent strand-independent λ-exonuclease biases in SNS-seq (Foulk et al. 2015; Petryk et al. 2016), SNS reads from non-replicating genomic DNA (LexoG0) were retrieved from SRA, accession SRP045284 (Foulk et al. 2015) and were used as a control during the peak calling step. Peaks (origins) were called using the MACS2 ‘callpeak’ function (Feng et al. 2012) with the following parameters: --gsize hs --bw 300 --qvalue 0.05 --mfold 5 50 and using LexoG0 uniquely mapped reads (in triplicate) as a control. Only origins detected in both HeLa SNS replicates were retained. Although using an algorithm designed for ChIP-seq peak calling might not be ideal for SNS-seq (Picard et al. 2014), we obtained a similar number of origins (~80,000) as reported using the optimized C++ SNS-scan detection algorithm (Picard et al. 2014). RT was assigned using the U2OS REPLI-seq data as described for the origins determined by EdU-seq.

### Hi-C data and topologically associating domain boundary definition

Topologically associating domain (TAD) maps at a 40-Kb resolution for 25 human cell lines were obtained from the 3D Genome Browser (Wang et al. 2018) in BED format (as detailed in **Table S1**). Most TAD maps were available in the hg19 genome build format, except those of 8 cell lines (GM12878, HCT116, HMEC, HUVEC, IMR90, KBM7, K562 and NHEK), which we downloaded in hg38 and converted to hg19 coordinates using the UCSC *liftOver* tool. All TAD maps were predicted from Hi-C data using the same previously described hidden Markov model (HMM) pipeline (Dixon et al. 2012). All intersections on the TAD maps BED files were performed using the *BEDtools* suite (Quinlan and Hall 2010). For each cell line we defined a set of 40 Kb boundaries, as described in McArthur and Capra (McArthur and Capra 2020). These were the 40-Kb windows surrounding (± 20 Kb) TAD start and stop coordinates. TAD boundary coordinates for each cell line are detailed in **Table S2**.

### Assignment of genic and intergenic TAD boundaries and genomic bins

Ensembl gene annotations (v99 for GRCh37/hg19) were used to create a list of all human genes and their position in the genome. Genomic bins or 40-Kb TAD boundary regions were classified as purely genic (Ge) if they mapped entirely within genes (forward, reverse or bidirectional direction of transcription), or purely intergenic (Ig) if they mapped entirely within intergenic sequences. If they contained both genic and intergenic sequences, they were classified as mixed (Ig-Ge). The analysis of the distribution of bins or boundaries in the genome considered the intergenic and mixed genic-intergenic bins as intergenic. A list of 2,176 human housekeeping genes, generated by mining thousands of RNA-seq datasets, was retrieved from the HRT Atlas v1.0 database (Hounkpe et al. 2020).

### Genomic features associated with TAD boundaries

ChIP-seq peaks from human cell line experiments for CTCF and members of the cohesin complex SMC3 and RAD21 were retrieved from ENCODE (Davis et al. 2018). For CTCF, we created a sorted BED file containing the intersection of peaks from experiments ENCSR000DNA, ENCSR000DLK, ENCSR000AMF, ENCSR000DPF, ENCSR000DMA, ENCSR000AKB, ENCSR315NAC, ENCSR000DWE, ENCSR000BNH, ENCSR000AKO, ENCSR000DKV, ENCSR000DZN, ENCSR000EFI, ENCSR000DYD, ENCSR000BPJ, ENCSR000DRZ, ENCSR240PRQ, ENCSR000DTO, ENCSR000BSE, ENCSR000EGM and ENCSR203QEB. For RAD21, we used the intersection of peaks from experiments ENCSR000DYE, ENCSR000BUC, ENCSR000BLD, ENCSR000FAD, ENCSR000BMY, ENCSR000BKV, ENCSR000EAC, ENCSR000EFJ, ENCSR000BSB and ENCSR000ECE. For SMC3, we used the intersection of peaks from experiments ENCSR481YWD, ENCSR000HPG, ENCSR000DZP and ENCSR000EGW. These experiments were selected based on the following search criteria: *assay type* ‘DNA binding’, *status* ‘released’, *genome assembly* ‘hg19’, *biosample* ‘cell line’ (selected from any of the 25 cell lines for which TAD maps were available, see **Table S1**), *file type* ‘BED NarrowPeak/BroadPeak’ and excluding any experiments with sample treatments (‘not perturbed’). The obtained files were intersected with the BED file containing the coordinates of all TAD boundaries to quantify ChIP-seq peak overlaps with each TAD boundary. Circular permutation tests were performed using the R package *regioneR* (Gel et al. 2016) to assess the significance of the genome-wide overlap between TAD boundaries and different genomic features, using *n* = 10,000 permutations. In addition, we tested if the association between TAD boundary positions and a given region set was dependent on their exact position by calculating local z-scores on 800-Kb windows around (±400 Kb) the exact positions of the observed overlap.

### Analysis of replication fork progression

To study fork restart and progression after hydroxyurea (HU) arrest and 90, 120 or 150 min post-release, we selected the early S-phase CN origins overlapping with TAD boundaries, as well as the early CN and Oi origins located within TADs. In particular, we selected start TAD boundaries which did not overlap with a preceding end boundary (similarly, end TAD boundaries non-overlapping with a subsequent start boundary). We calculated the average EdU-seq/release profiles ([0,1] range-normalized sigma values) over 600 Kb windows around these origins (±300 Kb). Computationally predicted CTCF binding sites were determined by scanning the FASTA files generated from these 600 Kb windows for the JASPAR core CTCF motif (MA0139.1) using the FIMO utility of the MEME software suite (Grant et al. 2011). Finally, the top-scoring (3rd quartile) significant CTCF motif matches (FDR < 0.05) were split by strand and their densities were calculated over the same 600 Kb windows using the generic kernel density estimation function in R.

### Data processing and availability

Datasets were downloaded and formatted using Unix shell scripting. Manuscript figures were created using Prism v8 (GraphPad), custom scripts in R v3.6.2 and previously published Perl v5.30.0 scripts (Macheret and Halazonetis 2018). No previously unreported algorithms were used to generate the results. Processed datasets generated in this study are available in the *TADs-replication* GitHub repository (https://github.com/EPL-repo/TADs-replication). The raw sequencing data used in this work is publicly available under the accessions detailed in **Table S1**. A summary of all non-base R packages used can be found in the table below:

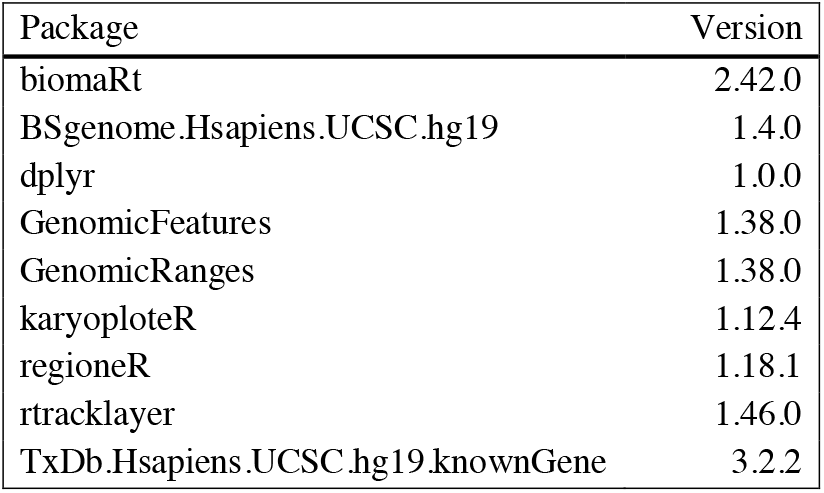

## SUPPLEMENTAL TABLES

**Supplemental Table 1.** Detailed sources and accession numbers for raw sequencing data.

**Supplemental Table 2. Coordinates of TAD start and end boundaries used in this study.**

## ACKNOWLEDGEMENTS

Research in M.T. laboratory is supported by Cancer Research UK, Medical Research Council and University of Oxford. This project has received funding from the European Union’s Horizon 2020 research and innovation programme under the Marie Skłodowska-Curie Individual Fellowship grant agreement No 886045 (‘BRCAstem’). E.P.L. is a recipient of a long-term EMBO non-stipendiary fellowship (ALTF 1044-2019).

---

**Figure S1.**
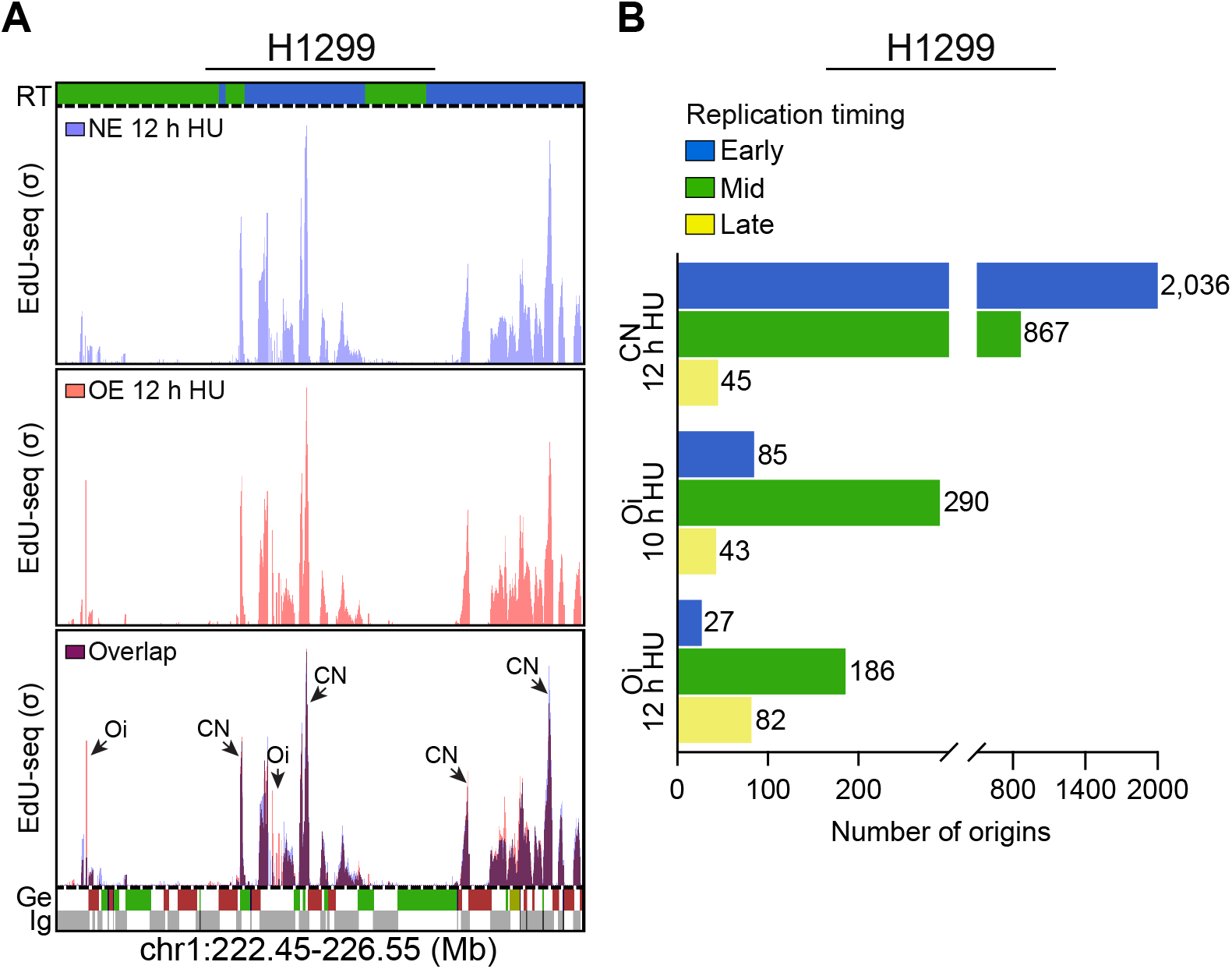
Quantification of localization and timing of replication origin firing in H1299 cells. **(A)** Representative genomic region of replication initiation profiles on chromosome 1 (chr1:222,450,000-226,550,000 bp). H1299 cells expressing normal levels of β-catenin (NE, purple) or overexpressing β-catenin (OE, red) were collected 12 h after mitotic shake-off. RT, replication timing profile of U2OS WT cells used as reference (blue, early; green, mid; yellow, late S-phase); σ (sigma), normalized number of sequence reads per 10 Kb-bin divided by its standard deviation; CN, constitutive origin; Oi, oncogene-induced origin; Ge, genic (green, forward direction of transcription; red, reverse; yellow, unspecified; blue, multiple genes within bin); iG, intergenic (grey). Bin resolution, 10 Kb. **(B)** Genome-wide quantification of CN (12 h after mitotic shake-off) and Oi (10 or 12 h after mitotic shake-off) according to their replication timing.

**Figure S2.**
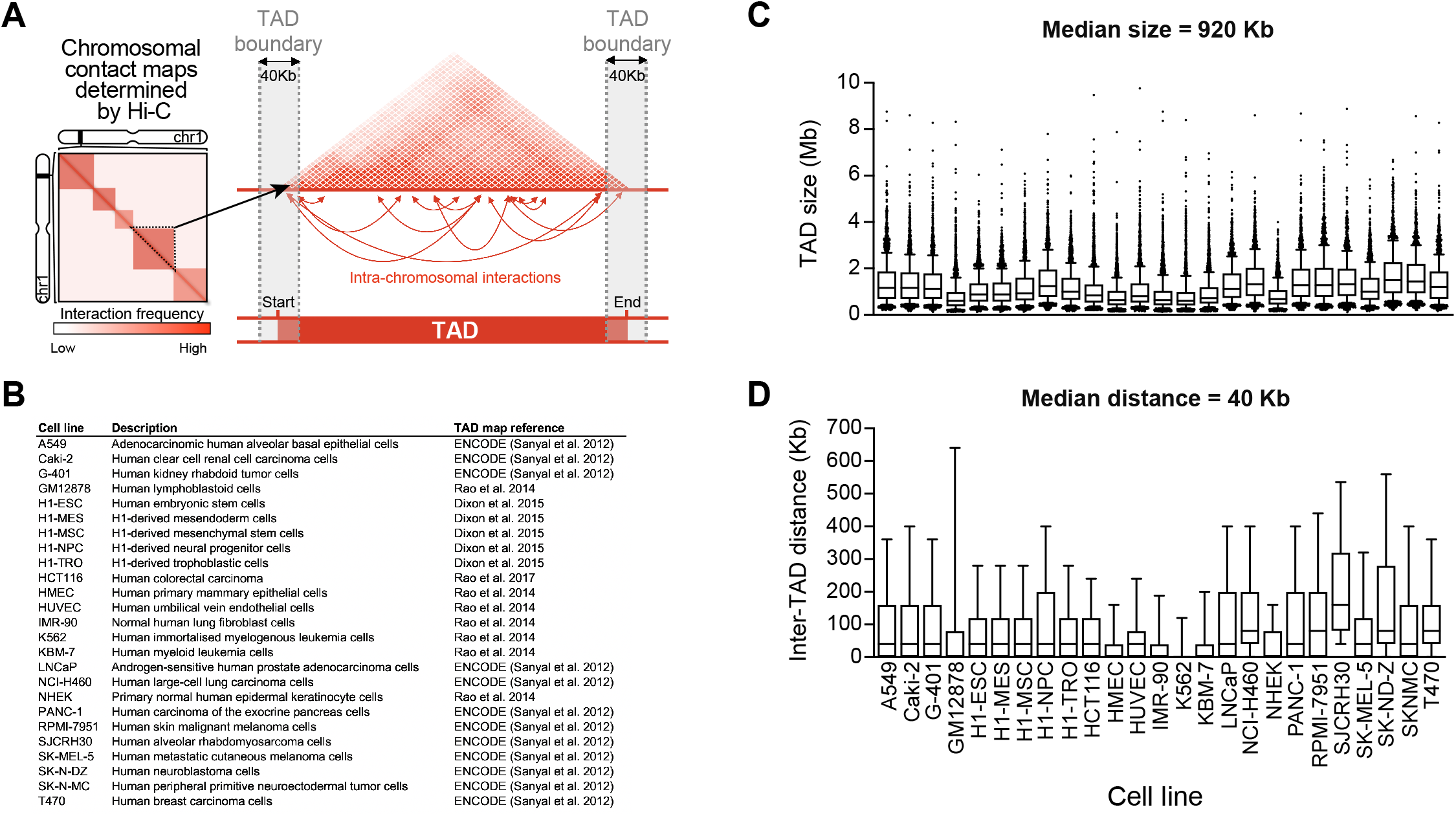
Defining the TAD maps used in this study. **(A)** Schematic representation of a TAD defined as self-interacting domains (loci within a TAD more frequently interact with one another than with loci outside of it) and visualised as triangles on Hi-C maps. Grey boxes represent TAD boundaries defined as regions bordering TADs and defined as 40-Kb windows surrounding (± 20 Kb) computationally determined TAD start and stop coordinates. **(B)** Twenty five cell lines and corresponding studies used to retrieve TAD from the 3D Genome Browser (Wang et al. 2018), at a 40-Kb resolution. **(C)** TAD size (in Mb) in each of the inspected cell lines. Whiskers represent the 10^th^ and 90^th^ percentiles. **(D)** Inter-TAD distance defined as distance (Kb) between the end coordinate of a given TAD and the start coordinate of the subsequent one, in each of the inspected cell lines. Whiskers represent the 10^th^ and 90^th^ percentiles.

**Figure S3.**
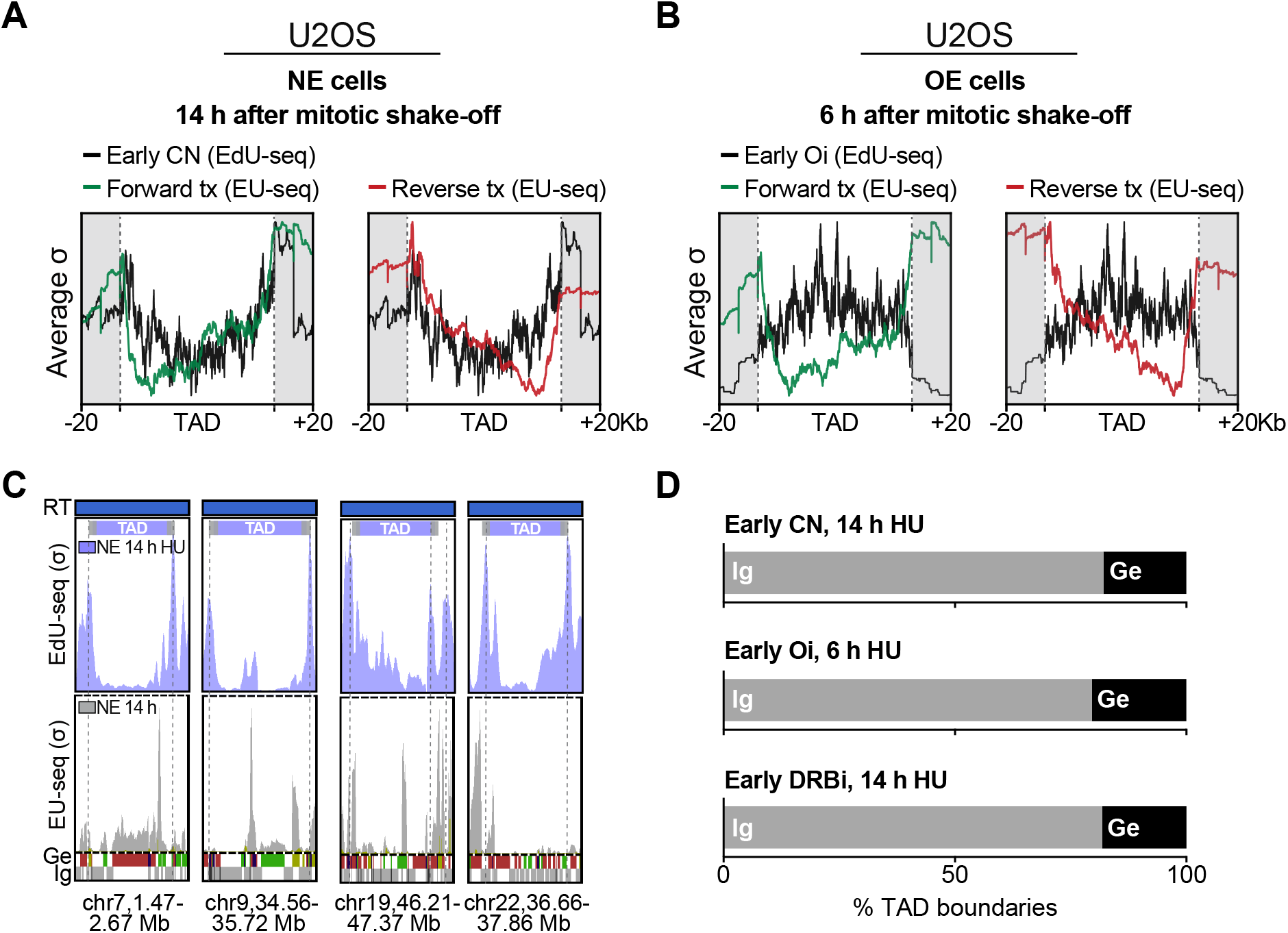
Transcription levels quantification along TADs. **(A-B)** Transcription profiles (based on average EU-seq σ values) are represented relative to TAD positions, with TAD boundaries (±20 Kb from TAD start/end points indicated by dotted lines) shadowed in grey. **(A)** Forward (green) and reverse (red) EU-seq average σ values in U2OS cells expressing normal levels of cyclin E (NE), collected 14 h after mitotic shake-off. **(B)** Forward (green) and reverse (red) EU-seq average σ values in U2OS cells overexpressing cyclin E (OE), collected 6 h after mitotic shake-off. EdU-seq signal for early **(A)** CN or **(B)** Oi origin firing (black) was superimposed to the EU-seq profiles. **(C)** Representative genomic regions showing early CN origin firing (EdU-seq) and transcriptional activity (EU-seq). U2OS cells expressing normal levels of cyclin E were treated with HU for 14 h. The grey dotted lines indicate peaks in EdU-seq signal. RT, replication timing (blue, early S-phase); σ, normalized number of sequence reads per 10 Kb-bin divided by its standard deviation; Ge, genic (green, forward direction of transcription; red, reverse; yellow, unspecified; blue, multiple genes within bin); iG, intergenic (grey); bin resolution, 10 Kb. **(D)** Percentage of intergenic (grey) and genic (black) TAD boundary regions overlapping with CN, constitutive origins; Oi, oncogene-induced origins; or DRBi, origins induced by DRB treatment of G1 cells (0-9 h after release form mitotic arrest).

**Figure S4.**
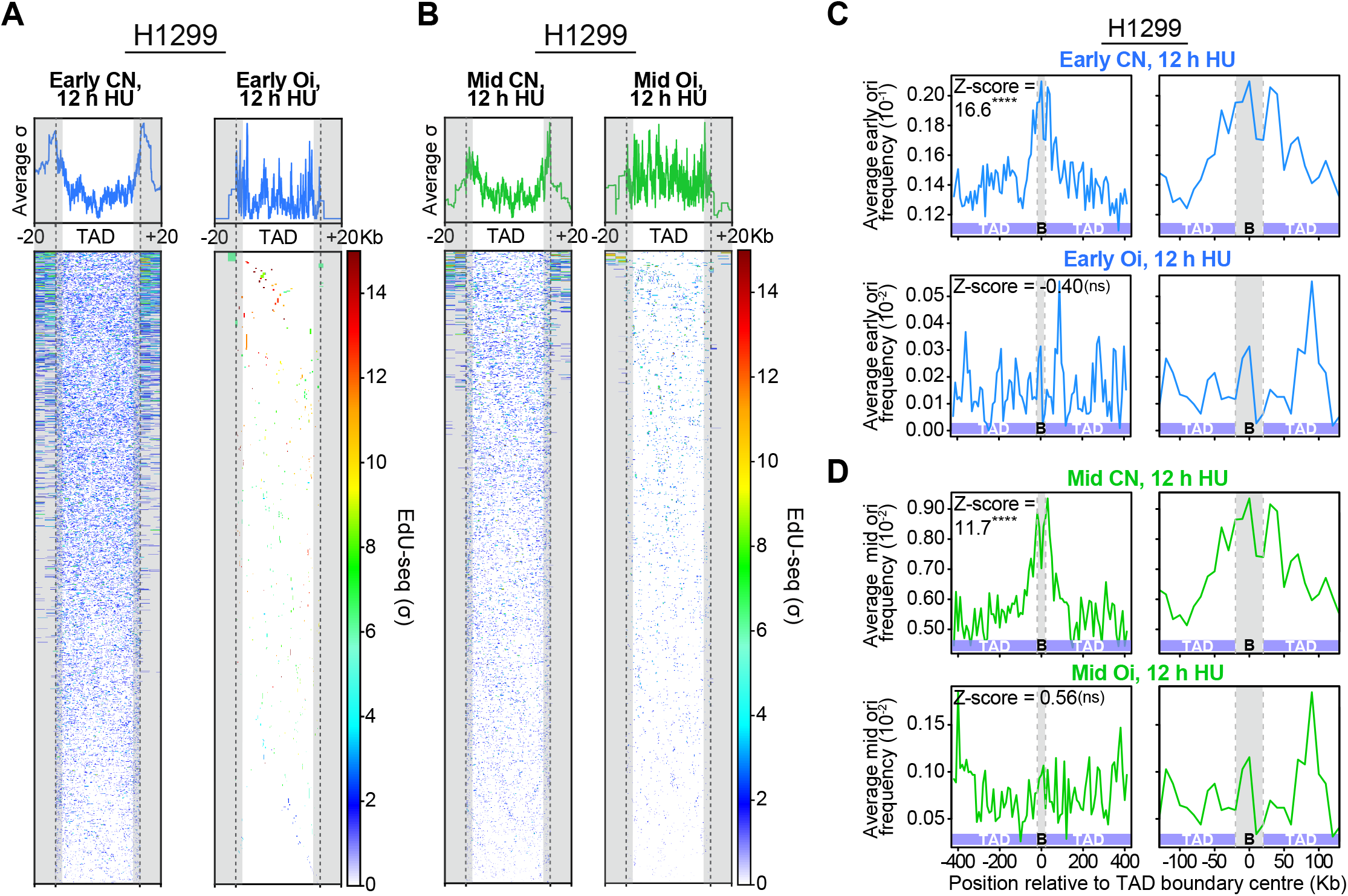
Distribution of replication origins firing in early or mid S-phase relative to TADs in H1299 cells. **(A-B)** Genome-wide early and mid S-phase replication origin firing profiles (based on average EdU-seq σ values) are represented relative to TAD positions, with TAD boundaries (±20 Kb from TAD start/end points indicated by dotted lines) shadowed in grey. Heatmaps represent EdU-seq σ values over each of the inspected TADs. **(A)** Early constitutive (CN) origins were mapped relative to the average position of *n* = 16,464 TADs having at least 1 early origin. Early oncogene (β-catenin)-induced (Oi) origins were mapped relative to the average position of *n* = 564 TADs having at least 1 early origin. **(B)** Mid CN origins were mapped relative to the average position of *n* = 11,718 TADs having at least 1 mid origin. Mid Oi origins were mapped relative to the average position of *n* = 2,758 TADs having at least 1 mid origin. TAD and TAD boundary mapping was based on average data obtained from 25 human cell lines shown in **Figure S2**. σ, normalized number of sequence reads per 10 Kb-bin divided by its standard deviation. **(C-D)** Frequency of replication origin firing within TAD boundaries and adjacent regions of ±400 Kb (left) or ±120 Kb (right). (C) Early S-phase origins. (D) Mid S-phase origins. Positive z-scores indicate that origins are enriched in TAD boundaries, whilst negative z-scores indicate that origins are depleted from TAD boundaries. ****, *P* < 1×10^−4^; ns, *P* > 0.05, permutations tests for association (*n* = 10,000 circular permutations). CN, constitutive origins; Oi, oncogene-induced origins. B, TAD boundary.

**Figure S5.**
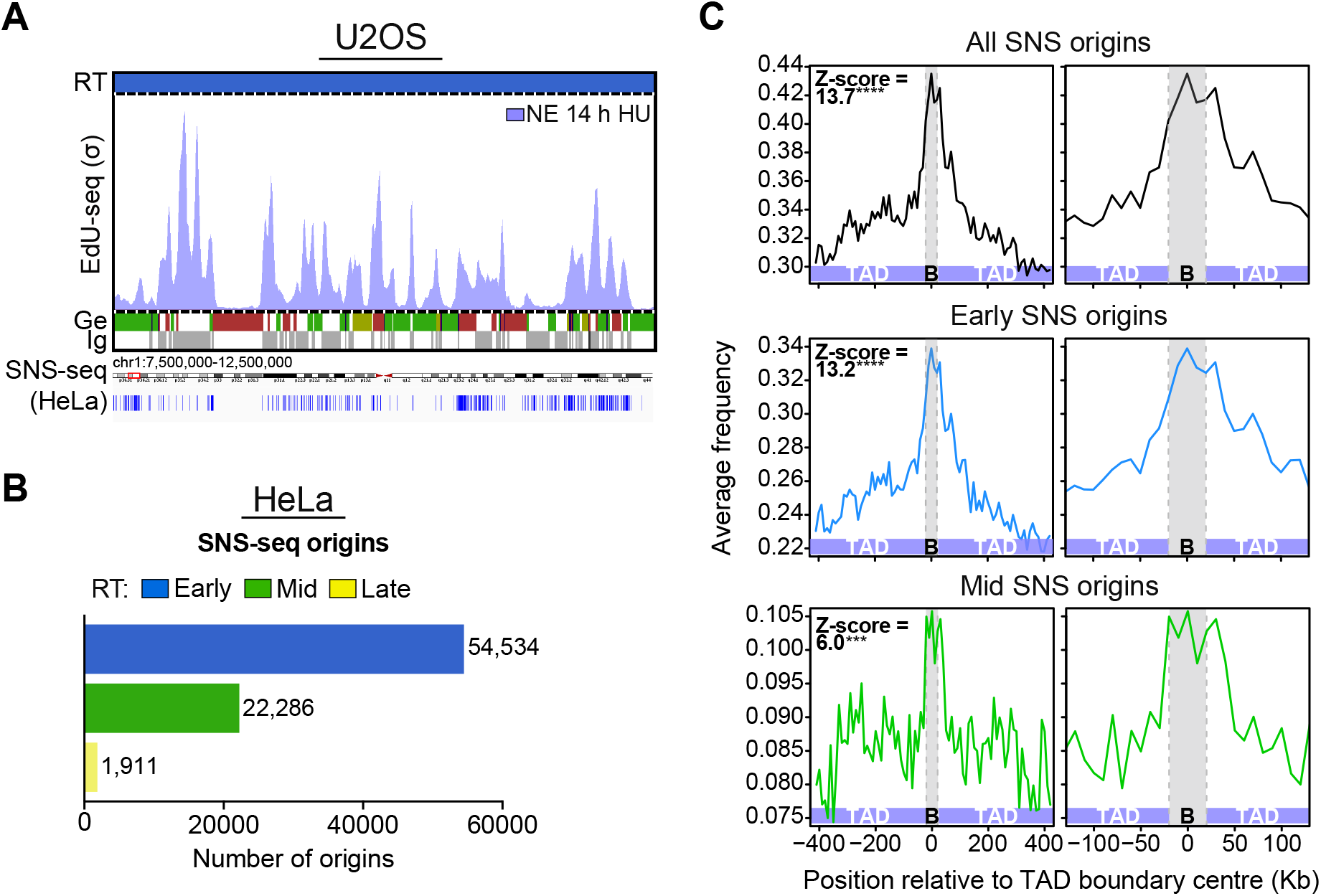
Distribution of replication origin firing determined by SNS-seq relative to TADs in HeLa cells. **(A)** Representative genomic region of replication initiation profiles determined by EdU-seq on chromosome 1 (chr1:7,500,000-12,500,000 bp). U2OS cells expressing normal levels of cyclin E (NE, light purple) were collected 14 h after mitotic shake-off. Discrete positions of origins determined by SNS-seq (HeLa cells) are represented on the bottom track. RT, replication timing profile of U2OS cells serving as reference (blue, early; green, mid; yellow, late S-phase); σ, normalized number of sequence reads per 10 Kb-bin divided by its standard deviation; Ge, genic (green, forward direction of transcription; red, reverse; yellow, unspecified; blue, multiple genes within bin); iG, intergenic (grey). Bin resolution, 10 Kb. **(B)** Genome-wide quantification of SNS-seq origins according to their replication timing. **(C)** Frequency of replication origin firing within TAD boundaries and adjacent regions of ±400 Kb (left) or ±120 Kb (right). Positive z-scores indicate that origins are enriched in TAD boundaries, whilst negative z-scores indicate that origins are depleted from TAD boundaries. ****, *P* < 1×10^−4^; ***, *P* < 1×10^−3^; ns, *P* > 0.05, permutations tests for association (*n* = 10,000 circular permutations). B, TAD boundary.

**Figure S6.**
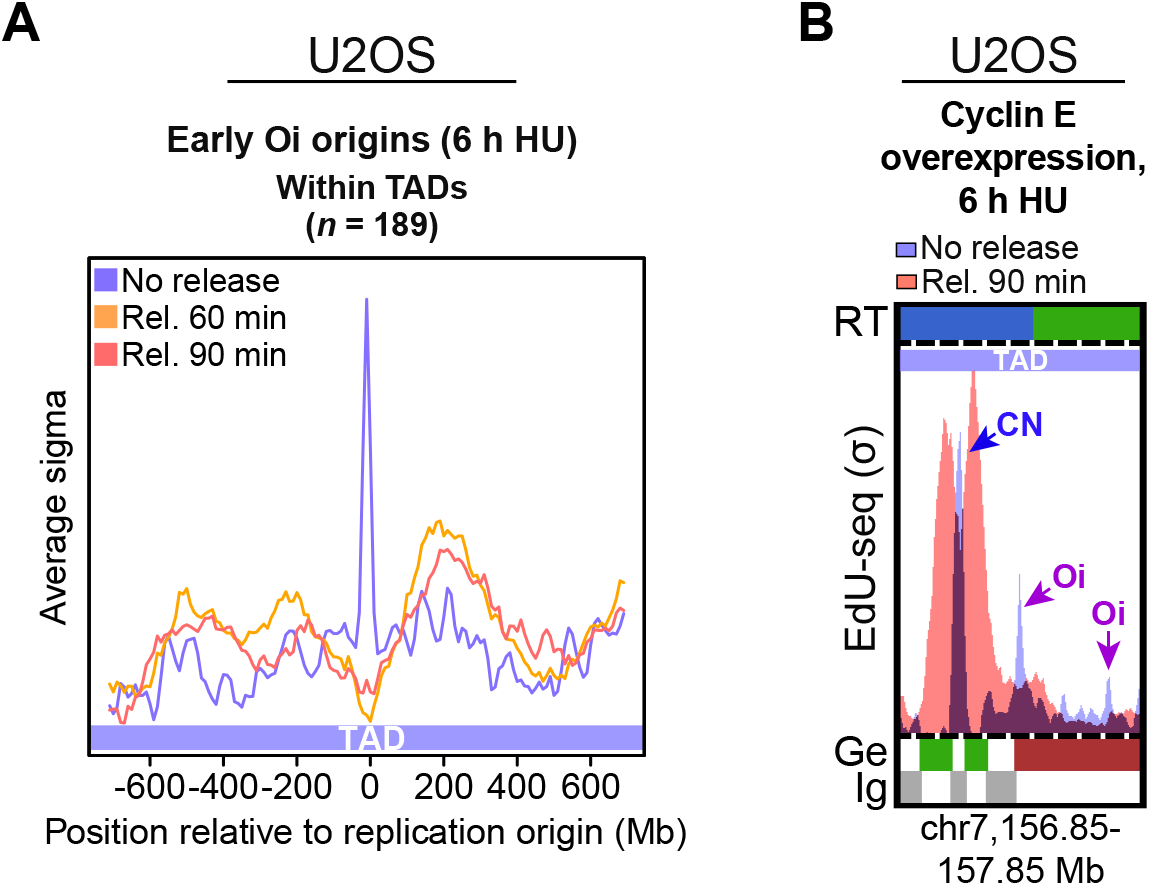
Forks initiating from Oi origins located within TADs collapse. **(A)** Average EdU-seq σ values showing early S-phase Oi origin firing in U2OS cells were treated with HU for 6 h (light blue) and fork progression in U2OS cells released from HU arrest for 60 min (orange) or 90 min (light red). B, TAD boundary. **(B)** Representative genomic regions showing Oi origin firing within TADs. U2OS cells were treated with HU for 6 h, followed by release for 90 min. The purple arrows indicate Oi origins and collapsed forks; the blue arrows indicate CN origins. RT, replication timing (blue, early; green, mid S-phase); σ, normalized number of sequence reads per 10 Kb-bin divided by its standard deviation; Ge, genic (green, forward direction of transcription; red, reverse; yellow, unspecified; blue, multiple genes within bin); iG, intergenic (grey); bin resolution, 10 Kb.

